# Dual-task Interference in a Simulated Driving Environment: Serial or Parallel Processing?

**DOI:** 10.1101/853119

**Authors:** Mojtaba Abbas-Zadeh, Gholam-Ali Hossein-Zadeh, Maryam Vaziri-Pashkam

**Affiliations:** School of Cognitive Sciences, Institute for Research in Fundamental Sciences, Niavaran, Tehran, Iran; School of Electrical and Computer Engineering, College of Engineering, University of Tehran, Tehran, Iran; Laboratory of Brain and Cognition, National Institute of Mental Health, Bethesda, Maryland, USA

**Keywords:** Dual-task interference, driving, drift diffusion model, task order predictability

## Abstract

When humans are required to perform two tasks concurrently, their performances decrease as the two tasks get closer together in time. This effect is known as dual-task interference. This limitation of the human brain could have lethal effects during demanding everyday tasks such as driving. Are the two tasks processed serially or in parallel during dual-task performance in naturalistic settings? Here, we investigated dual-task interference in a simulated driving environment and investigated the serial/parallel nature of processing during dual-task performance. Participants performed a lane change task on a desktop computer, along with an image discrimination task. We systematically varied the time difference between the onset of the two tasks (Stimulus Onset Asynchrony, SOA) and measured its effect on the amount of dual-task interference. Results showed that the reaction times (RTs) of two tasks in the dual-task condition were higher than those in the single-task condition. SOA influenced RTs of both tasks when they were presented second and the RTs of the image task when it was presented first. Manipulating the predictability of the order of the two tasks, we showed that unpredictability attenuated the effect of SOA by changing the order of the response to the two tasks. Next, using drift-diffusion modeling, we modeled the reaction time and choice of the subjects during dual-task performance in both predictable and unpredictable task order conditions. The modeling results indicated that performing two tasks concurrently, affects both the rate of evidence accumulation and the delays outside the evidence accumulation period, suggesting that the two tasks are performed in a partial-parallel manner. These results extend the findings of previous dual-task experiments to more naturalistic settings and deepen our understanding of the mechanisms of dual-task interference.

## 1. Introduction

One of the main characteristics of human cognition is its limited capacity. During driving, this limited capacity manifests itself when drivers attempt to simultaneously drive and perform a secondary task. Previous studies have shown that the presence of a secondary task increases reaction times (RTs) and reduces performance in driving (Blanco, Biever, Gallagher, & Dingus, 2006; Hancock, Lesch, & Simmons, 2003; Horrey & Wickens, 2004, 2006; Simmons, Caird, & Steel, 2017; Strayer & Fisher, 2016; Strayer & Johnston, 2001). To systematically investigate dual-task interference in non-driving tasks (Pashler, 1994a; Pashler & Johnston, 1989), the time interval between the onsets of the first and the second stimulus (henceforth referred to as the Stimulus Onset Asynchrony or SOA) has been systematically varied. It has been shown that when the SOA decreases, the reaction times increase, and the accuracies decrease. This performance decline as a function of SOA has been used as a measure of dual-task interference. Levy, Pashler, and Boer (2006) investigated this effect in a simulated driving environment. Participants performed a car following driving task while responding to a randomly presented visual or auditory stimulus using manual or verbal responses. The driving event was presented at different SOAs after the two-choice task. They showed that the reaction time of the driving task increased in the dual-task condition, while the reaction time of the visual/auditory task remained unchanged. Similar results were observed in another study (Hibberd, Jamson, & Carsten, 2013) using visual, auditory, and haptic stimuli for the secondary two-choice task. All these studies have presented the secondary task before the driving event. However, in natural driving conditions, a secondary task could appear either before or after the driving task. Here, we examined the dual-task interference in a simulated driving environment when the secondary task could appear at various time points relative to the driving task. Using the data collected in this naturalistic setting, we investigated if the two tasks are processed serially or in parallel during dual-task performance.

Several theories have been proposed to explain the dual-task interference; the two most influential of them are the “bottleneck theory” and the “central capacity sharing theory.” According to the bottleneck theory, dual-task interference appears when the two tasks rely on the same processor. In this model, this processor at any time can only be occupied by one of the two tasks (Pashler, 1994a). When the first task is being processed, the second task must wait for the first one to be finished so that the processor is released. Dividing each task into three stages of 1) perceptual, 2) response selection or decision, and (3) motor execution, the bottleneck model proposes that the stimulus perception and the motor execution stages could be performed in parallel, while the decision stage is the bottleneck that could only process the two tasks in a serial manner (McCann & Johnston, 1992; Sigman & Dehaene, 2008). Many studies have proposed evidence in favor of the bottleneck model (Pashler, 1994b; Pashler & Johnston, 1989; Ruthruff, Pashler, & Klaassen, 2001; Sigman & Dehaene, 2005). This theory predicts that the dual-task interference only affects the RT of the second task and has no effect on the response of the first task because the first task is processed by the decision stage first and postpones the processing of the second task (Pashler, 1994a).

On the other hand, the central capacity-sharing model suggests that the limitation in the processing capacity is the main reason for dual-task interference. Unlike the bottleneck theory that assumes serial processing of the two tasks, this theory suggests that in the dual-task conditions, all three stages of perceptual, decision and motor execution could process the two tasks in parallel (Duncan, 1980; Kahneman, 1973; McLeod, 1977; Posner & Boies, 1971). In this model, only the decision process is limited in capacity, while there are no resource limitations for the perceptual and motor execution stages (Michael Tombu & Jolicœur, 2003). This model predicts that dual-task interference affects the RT of both the first and the second tasks and that the size of this reaction time change depends on the size of the sharing portion. Several studies have provided evidence in favor of the capacity sharing model. Some have observed a robust effect of dual-task interference on the RT of both the first and the second tasks (Carrier & Pashler, 1995; Oriet, Tombu, & Jolicoeur, 2005; Sigman & Dehaene, 2006; Mike Tombu & Jolicœur, 2002; Zylberberg, Ouellette, Sigman, & Roelfsema, 2012). Others have shown that the process of decision making for the second task continues during the processing of the first task (Zylberberg et al., 2012), and factors such as task priority can influence the allocation of resources between the two tasks (McLeod, 1977; Michael Tombu & Jolicœur, 2003).

Evidence for or against dual-task theories are mostly gathered through simple tasks in artificial settings, and the predictions of these models have not been sufficiently tested in more naturalistic, real-world conditions. The use of a more naturalistic paradigm, such as driving, is crucial if the results of these theories are to be generalized to real-world everyday settings. The goal of this study is to measure the effect of SOA on the amount of dual-task interference in a simulated driving environment and to examine the validity of the capacity sharing and the bottleneck theories in more naturalistic settings.

As mentioned above, the main difference between the two dual-task theories is their predictions for the decision process of two tasks. One way to test the validity of the two interpretations of the dual-task interference is to precisely model the different stages of processing during the dual-task interference. A Drift diffusion model (DDM) could be used as a framework to model the processing of two-choice tasks (Ratcliff, 1978, 2015; Ratcliff & Rouder, 1998). This model assumes that during a two-choice decision task, evidence accumulates gradually to reach one of two decision thresholds corresponding to the two choices. The perceptual, motor and other non-decision related stages of task processing are modeled as the non-decision time in the DDM (Henceforth referred to as non-decision time, see Figure 2). The predictions of the bottleneck and the capacity sharing models can be restated within the framework of the DDM. The bottleneck model assumes that the two tasks are processed separately and sequentially, and that at shorter SOAs, the processing of the second task is delayed until the first task is completed. In other words, this model predicts that the rate of evidence accumulation (drift rate) for the two tasks is constant across SOAs, while there is a delay before the start of evidence accumulation for the second task that translates to increased non-decision time at shorter SOAs. On the other hand, the capacity sharing model suggests that the decision process for the two tasks can be operated concurrently, and the resources for decision making could be shared between the two tasks. This model predicts a decrease in the rate of evidence accumulation of the two tasks at shorter SOAs and a constant non-decision time across SOAs.

**Figure 1.**
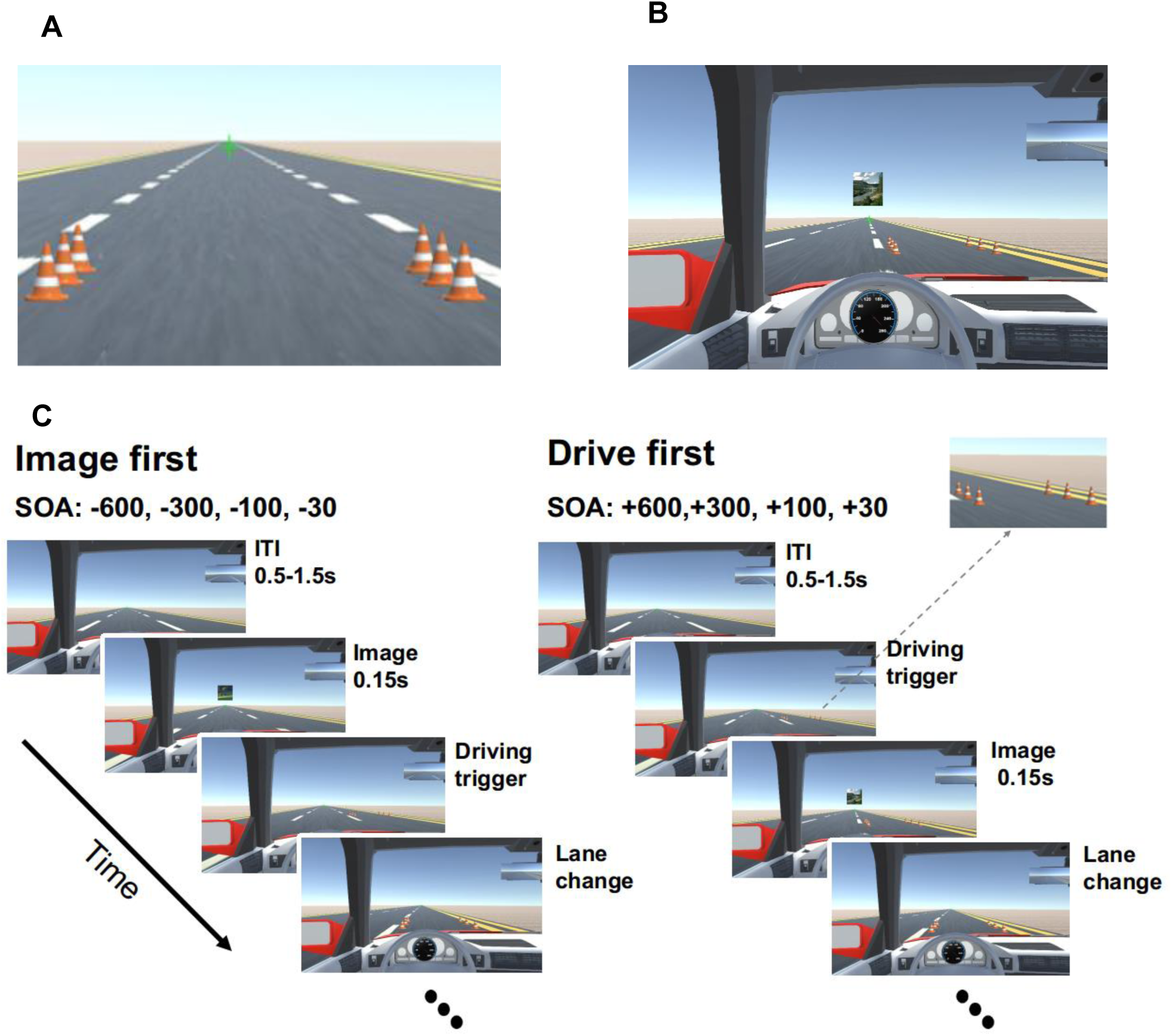
Dual-task paradigm. **(A)** A sample display showing the driving stimulus consisting of two rows of traffic cones in the middle driving lane. The cones were randomly presented in each lane, and participants had to drive through them without collision. **(B)** A sample display showing an image presented above the fixation point. Participants determined if the image was a face or a scene. **(C)** The sequence of events for a sample trial in which the image task was presented first (left), and another in which the driving task was presented first (right). The inter-trial interval (ITI) varied between 0.5 to 1.5 s. The image lasted for 150ms, and the cones were presented 30, 100, 300, or 600 before or after the image. Participants had to perform a lane change immediately after the appearance of the cones, and an image discrimination task immediately after the presentation of the image.

**Figure 2.**
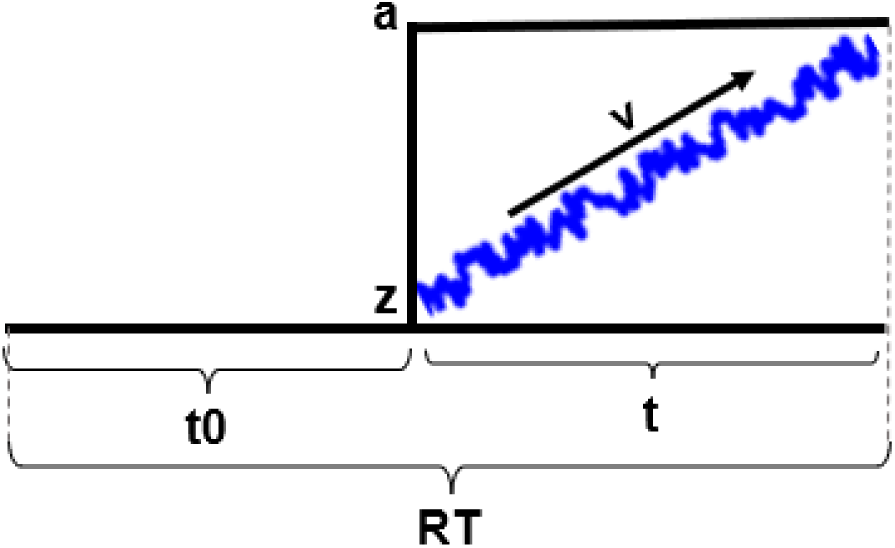
Schematic of the drift-diffusion process showing the DDM parameters. *t0* and *t* represent non-decision and decision times, *z* denotes the starting point of the decision process, *a* denotes the decision threshold, and RT denotes the reaction time.

DDMs have been used in previous investigations of dual-task interference. Zylberberg et al. (2012) using a dual-task paradigm with simple artificial tasks, have suggested that at shorter SOAs, the processing of the second task is not halted by the first task and that the evidence accumulation for the two tasks could happen concurrently. This is while the evidence accumulation for the second task decreases due to the interference from the first task. They suggested partial-parallel processing of the tasks during dual-task performance. In the current study, we aimed to extend these findings to a naturalistic setting and investigate the nature of dual-task interference in our simulated driving environment. To do this we explored the effect of SOA on driving performance and used a DDM to investigate if the driving and the secondary task are performed serially (central bottleneck theory) or in parallel (capacity sharing theory). In addition, we investigated if unpredictability of the order of the two tasks has an effect on driving behavior and the parameters of drift diffusion model.

In most dual-task studies, the order of the presentation of the tasks has been kept fixed and predictable, and participants were explicitly instructed to perform the two tasks according to the order of the presentation. In contrast, task order is often random and unpredictable in real-world situations. One open question is whether the order of the response to the two tasks during driving is specified based on a first-come, first-served basis, in which the order of the presentation determines the order of response, or this order is determined by a higher-order control mechanism.

In dual-task studies with simple designs (Sigman & Dehaene, 2005) in which the presentation order of the tasks is kept constant, and participants are often instructed to respond to the two tasks based on the presentation order, the first-come, first-served principle usually applies. However, recent studies which have made the order of the presentation of the two tasks unpredictable and have imposed no constraints for responding to the tasks according to the presentation order, support a higher-order control mechanism for managing the timing of the response to the two tasks (Fernández, Leonhard, Rolke, & Ulrich, 2011; Huestegge & Koch, 2010; Leonhard, 2011; Sigman & Dehaene, 2006; Szameitat, Lepsien, Von Cramon, Sterr, & Schubert, 2006). These studies have shown that increasing the perceptual difficulty of one of the tasks, such as degrading the stimulus, causes that task to be performed second (Sigman and Dehaene (2006); Strobach, Hendrich, Kübler, Müller, and Schubert (2018) but see also Leonhard, 2011 for evidence on the contrary). Similarly, when the difficulty of the decision (Fernández et al., 2011) or motor execution stages (Ruiz Fernández, Leonhard, Lachmair, Ulrich, & Rolke, 2013) is increased, participants are more likely to respond to that task later. These studies suggest that participants optimize the response order to decrease the total reaction time in dual-task conditions (Miller, Ulrich, & Rolke, 2009). All these studies have used simple artificial tasks rather than real-world naturalistic ones. It is still an open question if a higher-order control mechanism contributes to the response order in a naturalistic setting, such as a simulated driving environment. The second goal of this study is to measure the effect of task order predictability (OP) on the RTs and response order of the two tasks in naturalistic settings to investigate the possible involvement of higher-order control mechanisms in determining the response order during simulated driving. We will also explore the effect of OP on the parameters of the DDM.

In sum, we aim to investigate the effect of dual-task, SOA, and OP on subjects’ performance in a simulated driving environment and model the results using a DDM. The paradigm will consist of a lane change task and an image discrimination task. The image task will be randomly presented at different times (SOAs) relative to the lane change task. Participants will either perform both tasks or focus on one of the two tasks in separate blocks. We will investigate the effect of SOA and the OP of the two tasks on the amount of dual-task interference. Using a DDM, we will investigate whether the two tasks are processed in parallel or serially and how the OP influences the processing of the two tasks. Our investigation will help us achieve a deeper understanding of the mechanisms of dual-task interference in naturalistic settings, and could have implications for improving driving performance and reducing fatal accidents.

## 2. Material and Methods

### 2.1. Participants

Twenty healthy, right-handed adults (11 females), aged 20-30, participated in the study. All participants had normal or corrected to normal vision. Additionally, all participants were not expert video game players, as defined by having less than 2 hours of video-game usage per month in the past two years. All participants gave informed consent and were compensated for their participation.

### 2.2. Stimuli and Procedure

The dual-task paradigm consisted of a lane change driving task and an image discrimination task. The driving environment was designed in the Unity 3D game engine. Participants sat at a distance of 50 cm from a 22” LG monitor with a refresh rate of 60 Hz and resolution of 1920 × 1080 and responded to the tasks using a computer keyboard.

The driving environment consisted of a three-lane, desert road, without left/right turns or inclining/declining hills. Driving stimuli, composed of two rows of traffic cones (three cones in each row, see Figure 1), were presented on the two sides of one of the lanes in each trial, and the participants had to immediately redirect the car to the lane with the cones and pass through the cones. The space between the two rows of cones was such that the car could easily pass through them without collision. The cones were always presented in the lanes immediately to the left or immediately to the right of the car’s lane so that the participants had to change only one lane per trial. The lane change was done gradually: the subject had to hold the corresponding key to direct the car in between the two rows of cones, and then release the key when the car was situated correctly. Any early or late key press or release would cause a collision with the cones and a performance loss in that trial. The fixation cross was jittered for 100 ms to provide online feedback in case of a collision with the traffic cones. The participants were instructed not to change lane before the cones appeared. Trials in which participants changed lane before the presentation of the cones were considered false and removed from the analysis. Using this method, we could divide a continuous driving task into individual trials with predetermined onset and ends. At the beginning of the block, participants speeded up to 80 km/h using the “up” arrow key with the middle finger of the right hand. During the block, the speed was kept constant, and the subject moved to the left or right lanes by pressing the left and right arrow keys using the index and middle fingers of their right hand, respectively. For the image task, a single image of either a scene or a face was presented for 150 milliseconds centered at two degrees eccentricity above the fixation cross (Figure 1). The size of the image was 2.5 degrees of visual angle. Participants pressed the “x” and “z” keys on the computer keyboard with the middle and index fingers of their left hand to determine whether the image was a face or a scene, respectively. The images were pseudo-randomly selected from a set of 864 images of scenes and 435 images of faces. If participants responded incorrectly, the green fixation cross turned red, and if they responded late, it turned orange for 100ms. The length of each trial was three seconds, and the inter-trial interval varied randomly from 0.5 to 1.5 seconds. For the first trial in each block, the onset of the trial was set to be two seconds after the beginning of the block. The end of the trial was set to when the rear end of the car reached the end of the set of traffic cones.

To investigate the effect of task order predictability (OP) on dual-task interference, the experiment was performed in two different conditions: 1) “Predictable” task order condition and 2) “Unpredictable” task order condition. In two experimental conditions, the two tasks were presented with eight possible SOAs (−600, −300, −100, −30, +30, +100, +300 and +600). In the negative SOAs, the image was presented first (image-first), and in the positive SOAs, the driving was presented first (drive-first, Figure 1). In the Predictable conditions, the order of the presentation was fixed, so that in two of the four blocks, the driving task was presented first, and in the other two, the image task was presented first. In the Unpredictable condition, the order of the presentation of the two tasks was not predictable in each trial. Trials with driving as the first task were interleaved with trials with the image as the first task. Before the start of each block, participants were informed about the type of the block.

In addition to the dual-task conditions, participants performed two single-task conditions: 1) single driving task, and 2) single image task. In the single-task conditions, both the driving stimulus and the images were presented, but the subject only responded to one of them, ignoring the other. In the single image condition, the driving was on autopilot, and subjects only responded to the images. In the single driving condition, participants performed the lane change task and ignored the images.

Participants were told to focus on the fixation cross at the center of the page and respond to each task as fast as possible. At the end of each block, participants were informed about their performance on each task as well as their total performance. The performance in the driving task was calculated as the percentage of trials in which the subject passed through the cones without collision. The performance in the image task was calculated as the percentage of correct identifications.

Participants completed four blocks of 64 trials for each dual-task condition and two blocks of 32 trials for each single-task condition. There was a one-minute interval between blocks and a five-minute break after finishing all the blocks in each condition. The order of the blocks was counterbalanced across subjects.

Before performing the main experiment, all participants performed a block of 20 trials for every single-task. If their performance was 80% or higher, they proceeded to the main experimental blocks. Otherwise, they repeated blocks of 40 trials for each task until they reached 80% performance. After the single-task training, participants performed the dual-task training block. The dual-task training was similar to the single-task training block, with the difference that if after 20 trials, the dual-task performance did not reach the 75% threshold, the training was repeated with blocks of 50 trials.

### 2.3. Data Analysis

Only the correct trials were used for the reaction time analysis. In the dual-task conditions, if the response to both tasks was correct, that trial was included in the analysis. The trials in which the reaction time to each of the tasks was less than 200 ms and higher than 1500 ms were excluded from the analysis (3.48% of the trials). The analysis of response time and accuracy was performed using repeated-measures ANOVA with a Greenhouse-Geisser correction when necessary. False Discovery Rate correction (Benjamini & Hochberg, 1995) was applied in all cases that multiple comparisons were performed.

We used a logistic regression model to examine the effect of SOA and OP on the order of the response of the two tasks. The probability that the driving task was responded to first was determined by the following formula:

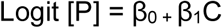

where P stands for the probability that the driving task was responded to first and C stands for SOAs. Parameters β_0_ and β_1_ were calculated for each subject. The model was fit separately on the data from the two dual-task conditions. A maximum likelihood estimation procedure was used for curve fitting.

### 2.4. Drift Diffusion Model fitting

To investigate if the two tasks were processed serially or in parallel, we used a drift-diffusion model (DDM) in which each trial was modeled as a combination of a non-decision time and a decision time consisting of a random drift towards decision bound (Figure 2). Model parameters consisted of: (1) parameter *z* denoting the starting point of the decision process; (2) parameter *a* denoting to the decision threshold; (3) parameter *v* representing the speed of information accumulation or drift rate; (4) parameter *t0* denoting the non-decision time pertaining to the combination of all other times in the trial excluding the drift-diffusion time. The DDM was implemented in the current study, by fitting the parameters *z, a, v*, and *t0*. We modified the DDM so that *z* and *a* were independent of SOA, and *v* and *t0* were dependent on SOA. Therefore, in the modified DDM, four values were fit for the parameter *v* and four values for the parameter *t0* corresponding to the four SOAs, one value for the parameter *a* and one value for the parameter *z* across all SOAs.

We used the Fast-dm package, developed by Voss and Voss (2007), for model fitting. Fast-dm is a package for fast drift-diffusion modeling. This package uses a partial differential equation method and a simplex routine to obtain the parameters of the DDM, and uses the calculated cumulative density function (CDF) of the predicted RTs to estimate the goodness of fit using a Kolmogorov–Smirnov (KS) function (Voss & Voss, 2008; Voss, Voss, & Lerche, 2015). The DDM was fit separately for each task (driving/image task) and each subject. We also calculated *R*^*2*^ values to further examine the goodness of fit of the model.

## 3. Results

### 3.1. Effect of Dual-task Interference on RTs

We first focused our analysis on the dual-task condition with the predictable task order and compared it with the single-task conditions (Figure 3). We ran four two-way repeated-measures ANOVAs with task condition (dual/single), and SOA as factors separately for the driving turn and the image discrimination and the drive-first and image-first task orders. Table 1 contains the details of the statistical results. Results showed a significant main effect of task condition with longer RTs in the dual-compared to the single-task condition in all cases (*ps* < 0.05). The effect of SOA was significant in all cases (*ps* < *0.05)* except for the driving turn RTs in the drive-first task order (*p = 0.15*). The interaction between task condition and SOA was also significant in all cases (*ps* < *0.05*). Further comparisons looking at the effect of SOA separately in the single- and dual-task conditions using one-way repeated-measures ANOVAs showed a significant effect of SOA on the RTs in both the single- and dual-task conditions in all cases (*ps* < *0.05*). The effect in the single-task condition might be driven by the presentation of the unattended stimulus that diverts participant’s attention from the main task (the presentation of the image in the case of the single driving task and the presentation of driving cones or the auto-pilot lane change event in the case of the single image task). The effect of SOA was also significant in the dual-task condition in all cases (*ps* < 0.05) except for the case of driving RT when it was presented first (*p* = 0.49). Consistent with previous studies of dual-task interference (Pashler & Johnston, 1989; Sigman & Dehaene, 2005; Mike Tombu & Jolicœur, 2002), when the image or the driving tasks were presented second, the RTs increased at shorter SOAs (*ps* < 0.01). Interestingly when the image was presented first, decreasing SOAs had an opposite effect, with shorter SOAs showing faster RTs (*p* < 0.001). These results have not been observed in previous dual-task studies and might be driven by participant’s urge to finish the image task sooner in order to reduce the interference on driving.

**Table 1.**
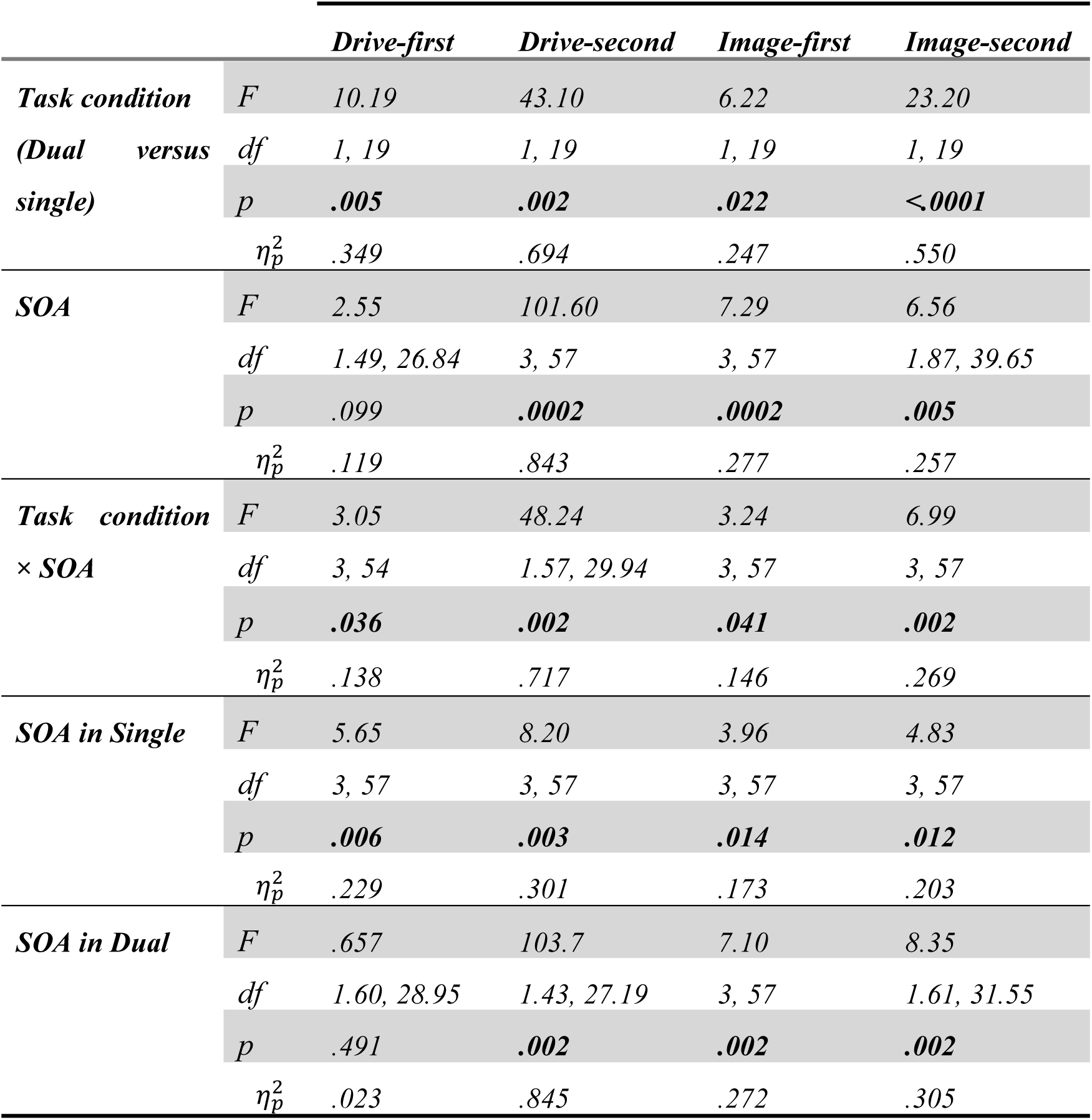
The first three rows show the results of two-way repeated-measure ANOVAs for the effect of task condition (dual vs. single), SOA, and the interaction between the two on RTs. The last two rows show the results of one-way repeated-measures ANOVAs for the effect of SOA on RTs separately in the single- and dual-task conditions. All p-values were corrected for multiple comparisons, and Greenhouse-Geisser correction was done when necessary.

**Figure 3.**
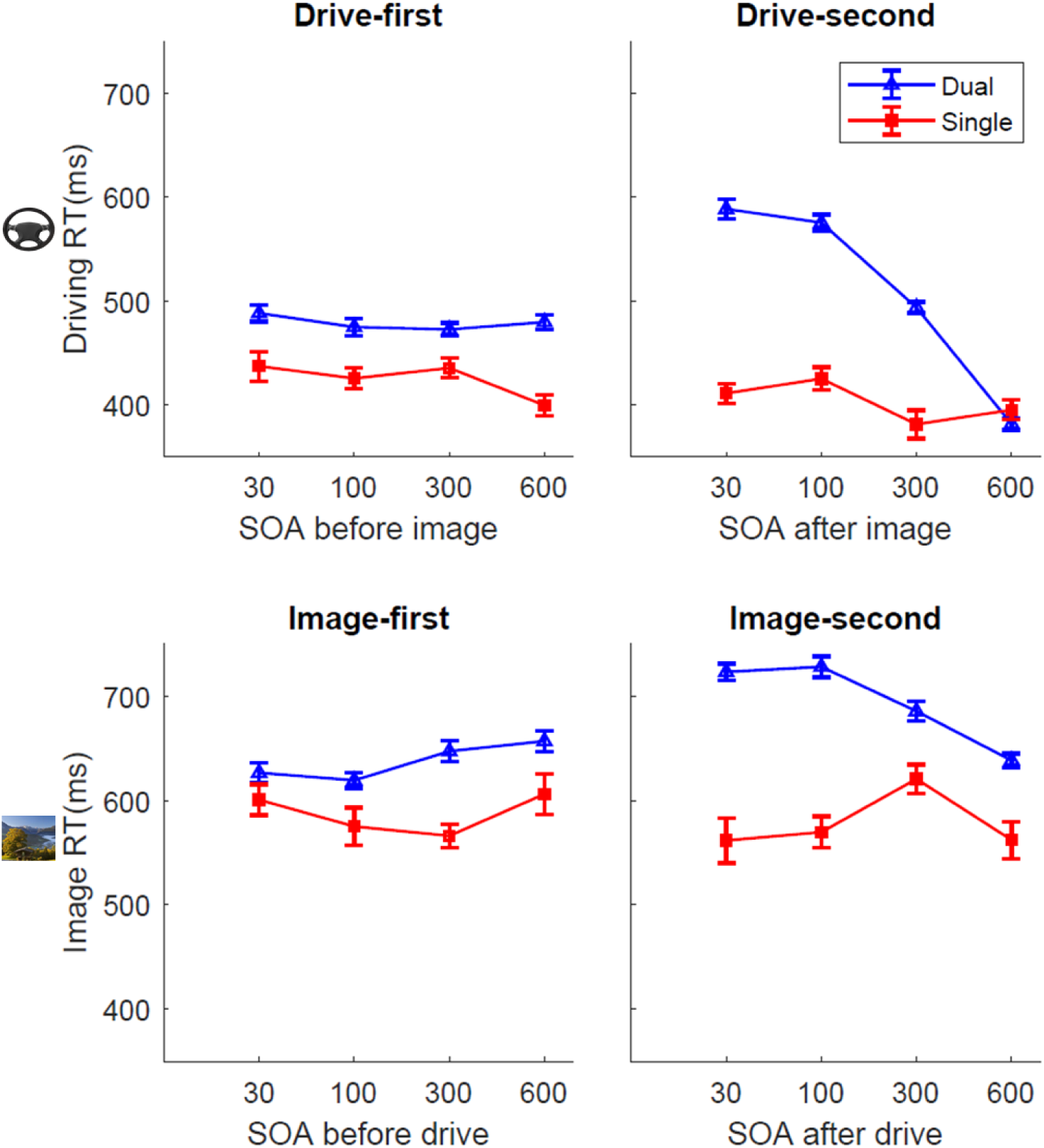
Effect of task condition (dual vs. single) and SOA on RTs. **(A, B)** These panels indicate the RTs for the driving turn in the drive-first and drive-second task orders, respectively, for the single-task (red) and the dual-task (blue) conditions. **(C, D)** These panels show the image discrimination RTs in the single (red) and the dual (blue) task conditions for the image-first and the image-second task orders, respectively.

In sum, our results show a clear effect of SOA on driving and image RTs. The presence of these strong effects allows us to use SOA as a factor for drift-diffusion modeling in the next section to investigate the nature of dual-task interference in our simulated driving set up.

### 3.2. Drift-diffusion Modeling of the Effect of Dual-task Interference on RTs

Drift diffusion modeling was used to investigate if a change in SOA affects the drift rate, non-decision time, or both. The model could account for most of the variance in the data (R^2^: Drive-first 0.78±0.03, Drive-second 0.94 ±0.02, Image-first 0.71 ±0.04 and Image-second 0.84±0.03) and the distribution of the RTs from the model fit was not significantly different from the original data in all subjects and all conditions (*ps* > 0.1).

Next, we investigated the effect of SOA on the two model parameters *v* and *t0*, corresponding to the drift rate and non-decision times. Serial processing of the two tasks would lead to an increase in the *t0* at shorter SOAs for the second task, while parallel processing of the two tasks would decrease the *v* for both tasks. Results showed that when either of the two tasks was presented second, *v* decreased and *t0* increased at shorter SOA (*ps* < 0.05, Figure 4B, D, F & H). No significant change in *v* or *t0* was observed when driving was presented first (*p* > 0.05, Figure 4A & E) and a decrease in both *t0 v* was observed at shorter SOAs when the image was presented first (*ps* < 0.05, Figure 4C & G). The details of statistical tests are shown in table 3. These results suggest that the two tasks are neither processed in a strictly parallel nor a strictly serial manner, as a change in the non-decision time is always accompanied by a change in the drift rate.

**Table 3.** One-way repeated-measures ANOVAs for the effect of SOA on *v* and *t0.* All p-values were corrected for multiple comparisons and Greenhouse-Geisser correction was done when necessary.

|  |  | *Drive-first* |  | *Drive-second* |  | *Image-first* |  | *Image-second* |  |
| --- | --- | --- | --- | --- | --- | --- | --- | --- | --- |
| | | $v$ | $t0$ | $v$ | $t0$ | $v$ | $t0$ | $v$ | $t0$ |
| **SOA** | *F* | 1.59 | 3.46 | 15.03 | 64.42 | 3.95 | 7.96 | 6.67 | 12.20 |
|  | *df* | 1.87, | 1.60, | 3, 57 | 1.59, | 3, 57 | 1.35, | 3, 57 | 1.86, |
|  |  | 35.53 | 30.43 |  | 30.37 |  | 25.80 |  | 35.36 |
|  | *p* | .217 | .663 | .002 | .001 | .022 | .006 | .002 | .001 |
| | $\eta_p^2$ | .078 | .018 | .442 | .772 | .172 | .295 | .296 | .391 |

**Figure 4.**
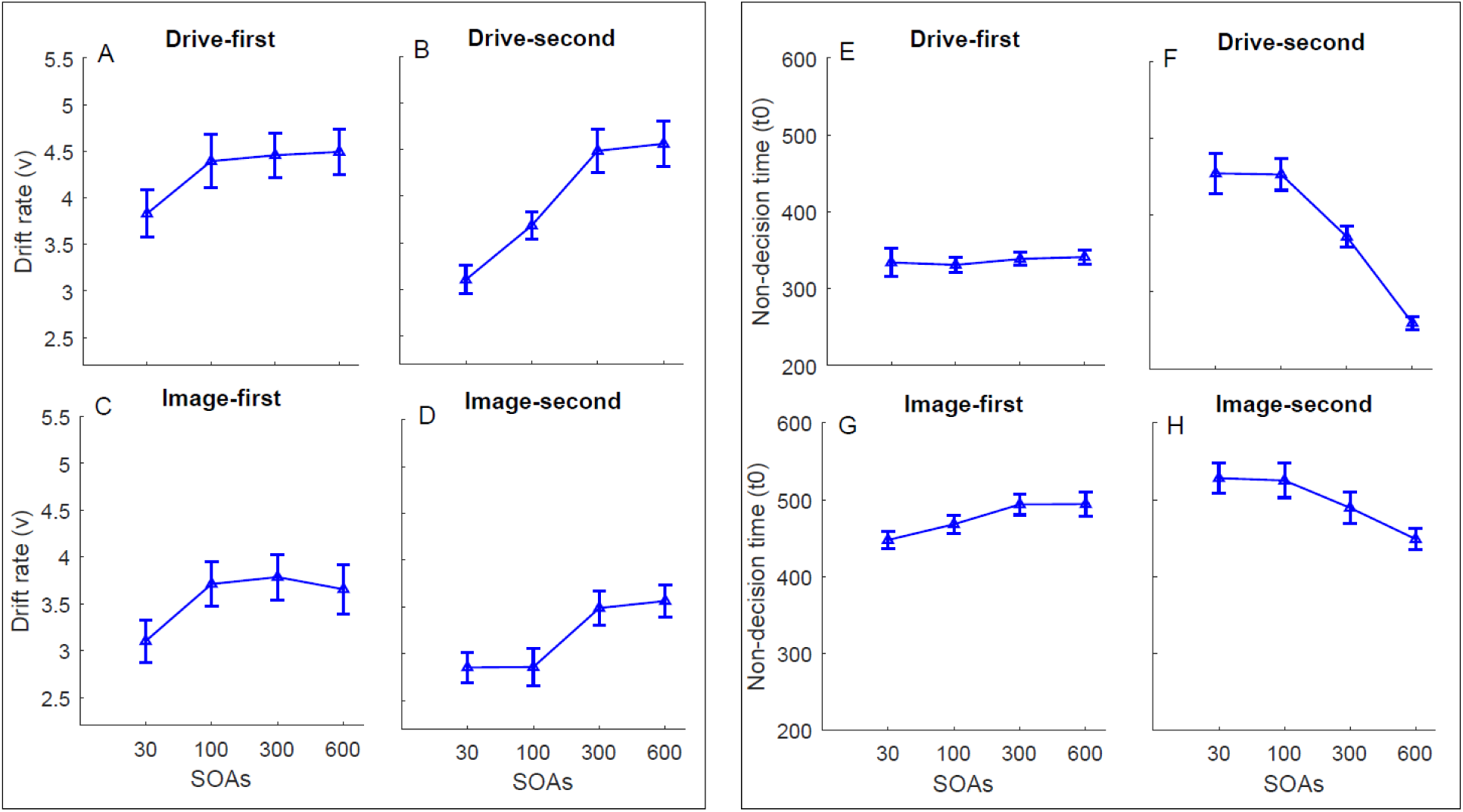
Effect of SOA on drift rates (*v*) and non-decision times (*t0*). Panels A-D on the left show the effect of SOA on the drift rate (v) for driving turn in the drive-first **(A)** and drive-second **(B)** conditions and that for the image discrimination in the image-first **(C)** and image-second **(D)** conditions. Panels E-H on the right show the effect of SOA on non-decision time (t0) for driving task in the drive-first **(E)** and drive-second **(F)** conditions and that for the image task in the image-first **(G)** and image-second **(H)** conditions.

### 3.3. Effect of Task Order Predictability on RTs

To investigate the effect of task order predictability (OP) on the RTs during dual-task performance, we compared the main dual-task condition in which the task orders were predictable (i.e., the two task orders were presented in separate blocks) to a condition in which the task orders were unpredictable and varied randomly from trial to trial within a block. We ran four two-way repeated-measures ANOVAs with task condition (predictable/unpredictable) and SOA as the two factors, separately for the driving turn and the image discrimination, and the drive-first and image-first task orders. The details of the statistical tests are summarized in table 4. The effects of OP, SOA, and their interaction on RTs were significant in both drive-first and drive-second conditions (*p* < 0.05; Figure 5A & B). When the image was presented first (Figure 5C), OP had a marginally significant effect on mean image RTs (*p* = 0.07), and the interaction between OP and SOA was significant (*p* > 0.03). When the image was presented second (Figure 5D), the effect of OP on RTs (*p* = 0.052), and the interaction between OP and SOA on RTs (*p* = 0.057) were marginally significant.

**Table 4.** Results of two-way repeated-measures ANOVAs for the effect of OP and SOA on RTs and one-way repeated-measures ANOVAs for effect SOA on RTs in the unpredictable condition. All p-values were corrected for multiple comparison and Greenhouse-Geisser correction was done when necessary.

|  |  | *Drive-first* | *Drive-second* | *Image-first* | *Image-second* |
| --- | --- | --- | --- | --- | --- |
| **OP** | *F* | 10.21 | 11.81 | 3.53 | 4.29 |
|  | *df* | 1, 19 | 1, 19 | 1, 19 | 1, 19 |
|  | *p* | .005 | .012 | .076 | .069 |
| | $\eta_p^2$ | .350 | .383 | .157 | .184 |
| **SOA** | *F* | 2.83 | 88.89 | 3.35 | 7.55 |
|  | *df* | 1.48, 28.27 | 1.51, 28.82 | 3, 57 | 1.65, 31.50 |
|  | *p* | .089 | .0004 | .050 | .006 |
| | $\eta_p^2$ | .130 | .824 | .150 | .285 |
| **OP × SOA** | *F* | 5.02 | 7.06 | 3.37 | 3.07 |
|  | *df* | 3, 57 | 3, 57 | 3, 57 | 1.83, 34.82 |
|  | *p* | .003 | .004 | .041 | .057 |
| | $\eta_p^2$ | .215 | .271 | .151 | .139 |

**Figure 5.**
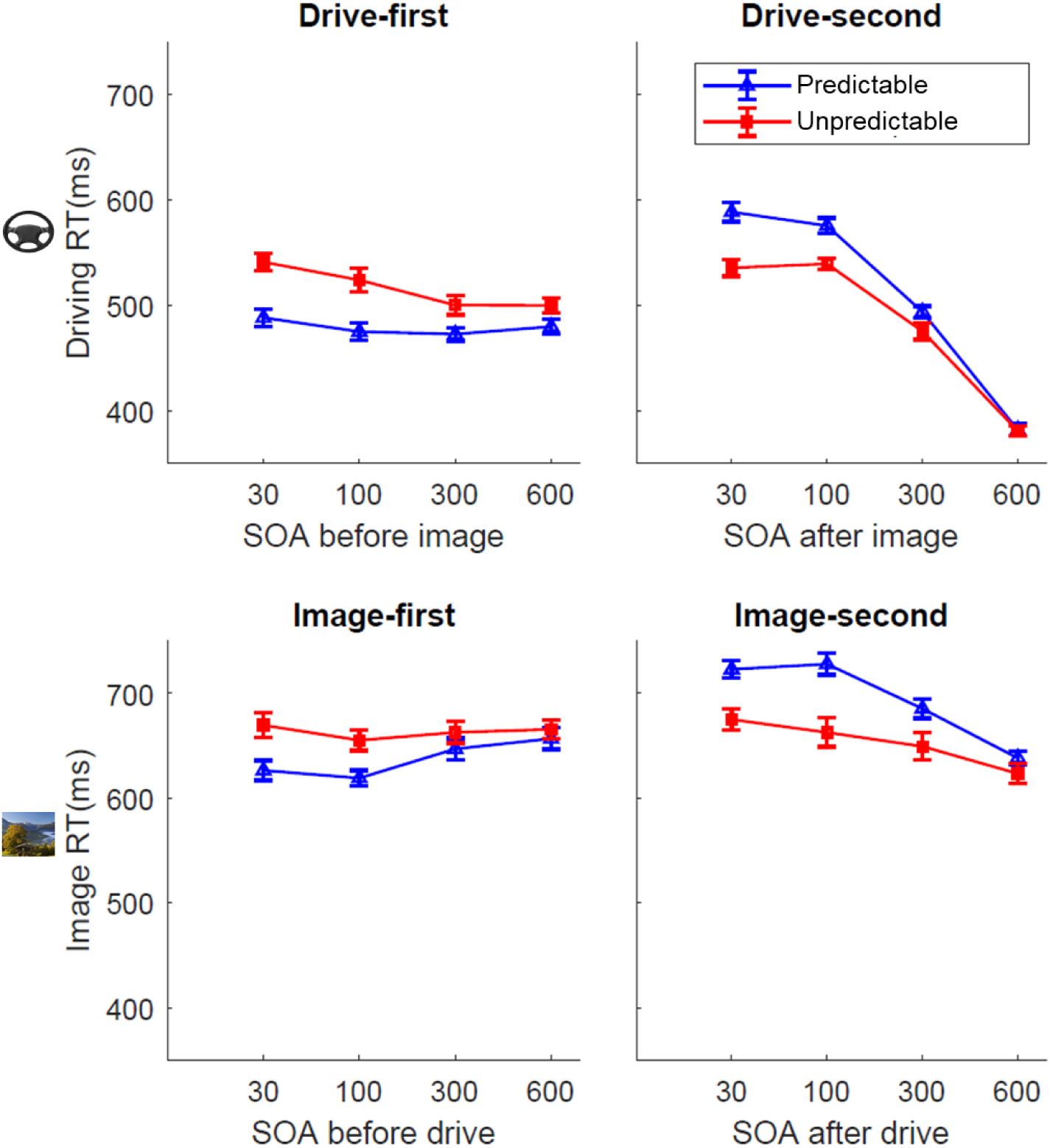
Effect of OP and SOA on RTs. The two top panels show the RTs for the driving turn in the drive-first **(A)** and drive-second **(B)** task orders for the predictable (blue) and the unpredictable (red) task order conditions. The two bottom panels show the RTs for the image discrimination task in the image-first **(C)** and image-second **(D)** task orders for the predictable (blue) and the unpredictable (red) task order conditions.

Furthermore, we investigated the effect of SOA separately in the unpredictable conditions using one-way repeated-measures ANOVAs (note that the effects for the predictable condition are already reported in the previous section). The results showed a significant effect of SOA on the RTs in all cases (*ps* < 0.05) except for when the image was presented first (*p* = 0.56).

In general, these results demonstrate that OP increases the mean RT of the first task and decreases the mean RT of the second task with the changes more pronounced when the tasks get closer together in time. These results show that unpredictability of the task order attenuates the effect of SOA on RTs for all cases except the drive-first RTs. We next investigated the possible origin of this attenuation effect.

### 3.4. Effect of Task Order Predictability on the Response Order

To investigate the effect of SOA and OP on the order of the response to the two tasks, we calculated the probability that the driving task was responded to first in each SOA and for each subject (Figure 6A) and fit a logistic regression model to these probability values. The model was fit separately for each of the two dual-task conditions and an intercept (β0 in the logistic model described in the methods) and a slope (β1 in the logistic model) was calculated for each condition and each subject. We also calculated the SOA value in which the probability of responding to the driving task first was 50% (T50). Then, to quantify the effect of OP on the response order, the model outputs and the T50 value across the two experimental conditions were submitted to a paired t-test. OP had no significant effect on the shift (β1) of the logistic function (*t* (1,19) = 0.323 *p* = 0.75, see Figure 6B). The slope of the logistic function (β1) was significantly influenced by OP (*t* (1,19) = 3.08, *p* = 0.006). Negative T50 values in both conditions show that participants had a general bias to respond to the driving task first (Figure 6C) but this bias was the same across the two conditions (*t* (1, 19) = .317, *p* = 0.75). At SOA = 0, in more than 60 percent of trials driving was responded to first. In sum, these results showed that OP changes the response order to the two tasks and has no effect on the bias in favor of the driving task.

### 3.5. Drift-diffusion Modeling of the Effect of Task Order Predictability on RTs

DDM was fit to the data from the predictable and unpredictable task order conditions, separately, and output model parameters were compared for the two conditions. The result of model fitting on the unpredictable task order condition showed that the model could account for most of the variance in the data (R^2^: Drive-first 0.70±0.04, Drive-second 0.96±0.01, Image-first 0.75±0.03 and Image-second 0.82±0.03) and the distribution of the RTs from the model fit was not significantly different from that of the original data in all subjects and all conditions (*ps* > 0.09). We ran two-way repeated-measures ANOVAs to investigate the effect of task condition (Predictable vs. Unpredictable) and SOA on the two parameters *t0* and *v*, separately for the two task orders, and the driving and the image discrimination tasks. The details of the statistical test are shown in table 4. The effect of OP on *v* was not significant in all cases (*ps* > .05, Figure 7A, B, C & D). This effect on *t0* was only significant in the drive-second (*p = 0*.012, Figure 7F) and marginally significant for image-second conditions (*p =* 0.052, Figure 7H) and was not significant in the drive-first and image-first conditions (*ps* > 0.05, Figure 7E & G). SOA had a significant effect on *v* and *t0* in all conditions (*ps* < 0.05), except when the driving task was presented first (*p* > .05, Figure 7A). The interaction of OP and SOA on *t0* was only significant for drive-second conditions (*p =* 0.003, Figure 7F). These results show that when either the image or the driving tasks were presented second, unpredictability changed the non-decision time of the tasks. Note that the analysis of the response order showed that in the unpredictable condition, the second task was more likely to be responded to first. The changes in the order of response could be tightly related to the decrease in the non-decision time of the second task.

**Figure 7.**
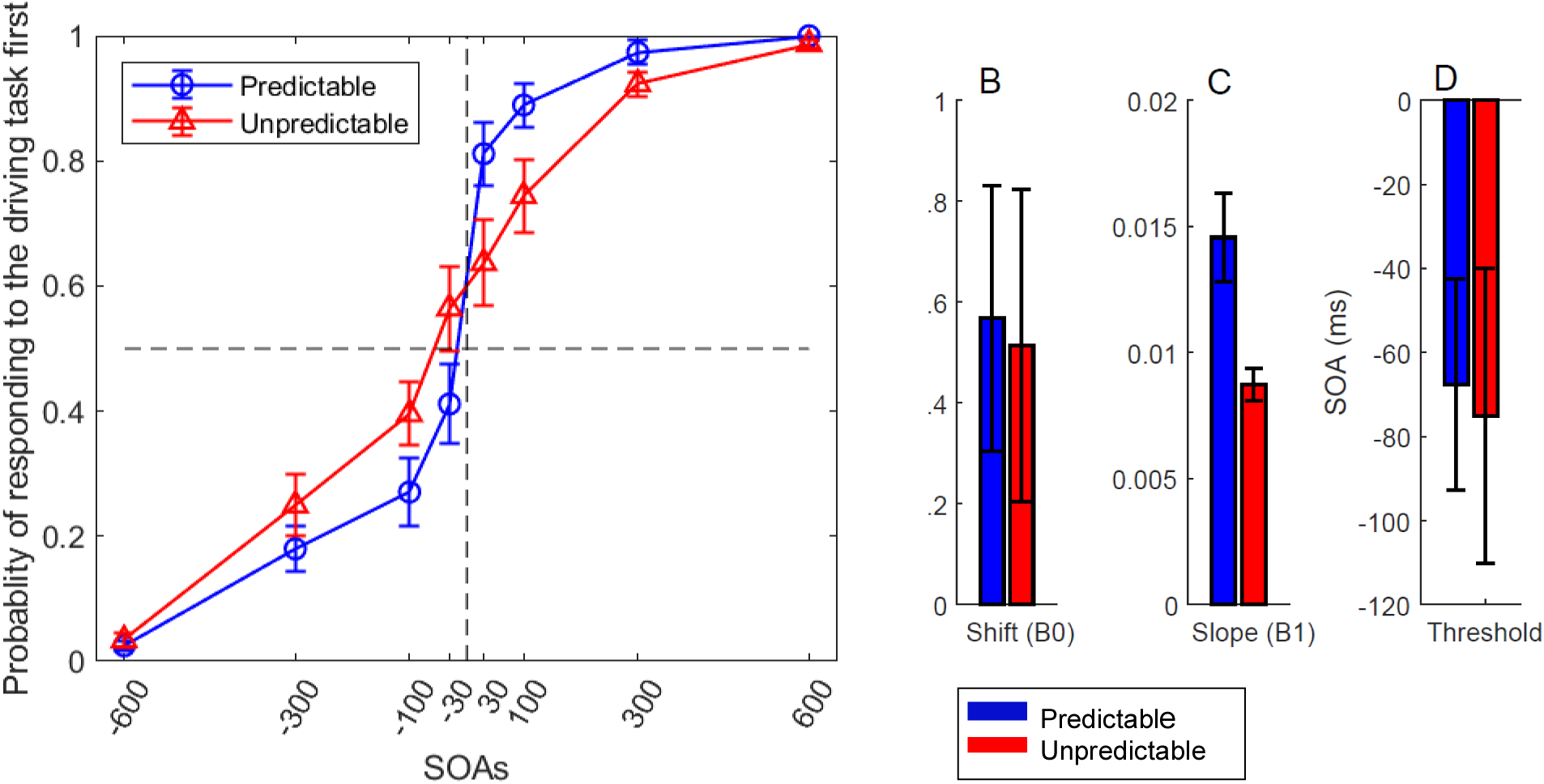
Effect of OP on the response order. Predictable and Unpredictable conditions are shown in blue and red colors, respectively. **(A)** The probability of first responding to the driving task plotted for the two task conditions. The curves are fit to the average data using a logistic regression function. **(B)** The shift of the logistic regression function (β0), **(C)** the slope of the logistic function (β1), and **(D)** The T50 (the SOA in which participants responded to the driving task first with 50% probability), for the predictable (blue) and unpredictable (red) conditions. The shift did not differ between the two conditions, but the slope was shallower in the unpredictable condition (*p* < 0.006). There was a general bias for responding to the driving task first in both conditions.

**Figure 7.**
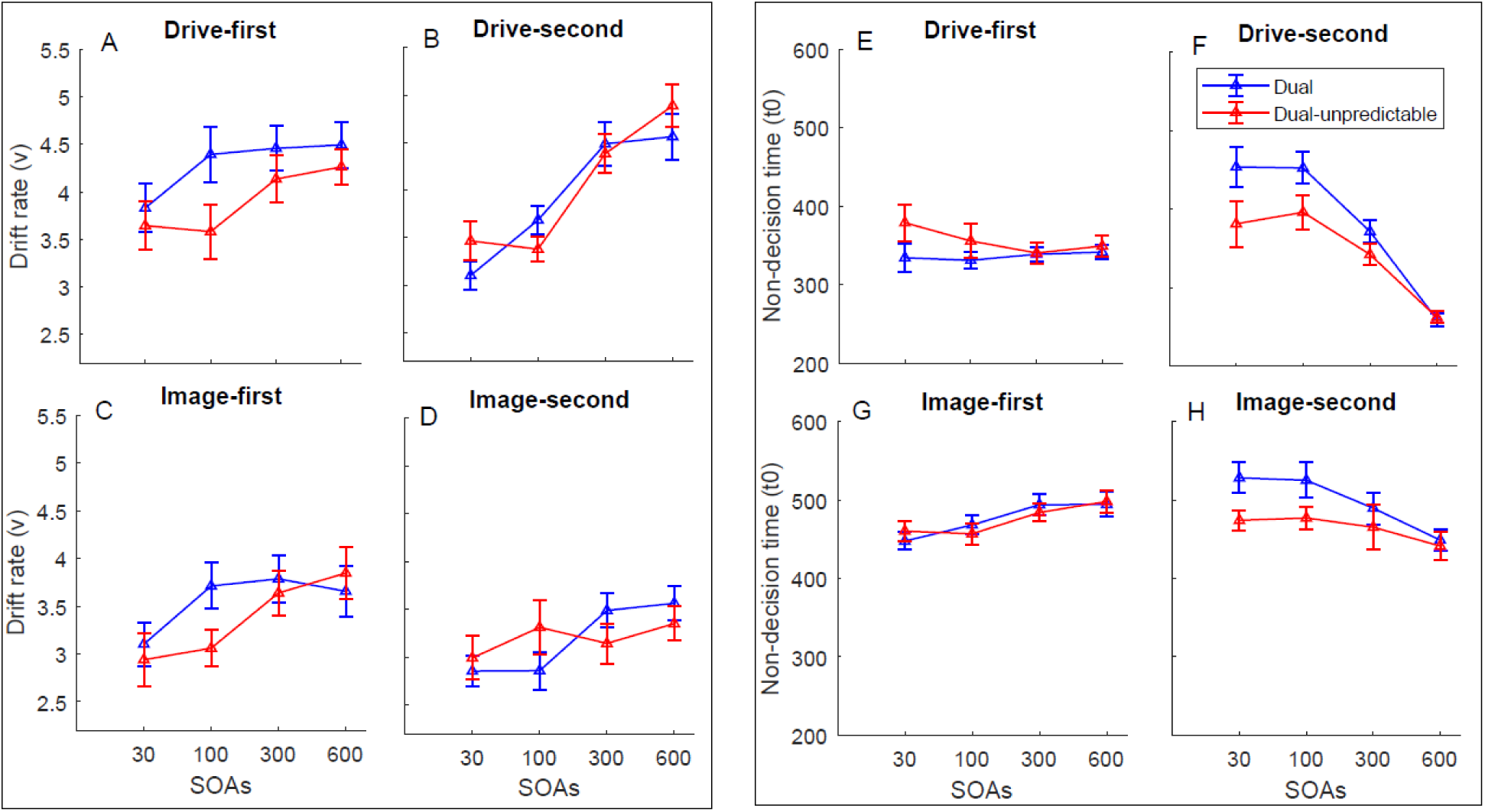
Effect of OP and SOA on drift rates (*v*) and non-decision times (*t0*). Blue lines and red lines show the predictable and unpredictable task orders, respectively. Panels A-D on the left show the effect of OP and SOA on the drift rate (v) for driving turn in the drive-first **(A)** and drive-second **(B)** conditions and that for the image discrimination in the image-first **(C)** and image-second **(D)** conditions. Panels E-H on the right show the effect of OP and SOA on non-decision time (t0) for driving task in the drive-first **(E)** and drive-second **(F)** conditions and that for the image task in the image-first **(G)** and image-second **(H)** conditions.

**Table 4.** Results of two-way repeated-measures ANOVAs for the effect of OP and SOA on *v* and *t0*. All p-values were corrected for multiple comparisons and Greenhouse-Geisser correction was done when necessary.

|  |  | *Drive-first* |  | *Drive-second* |  | *Image-first* |  | *Image-second* |  |
| --- | --- | --- | --- | --- | --- | --- | --- | --- | --- |
| | | $v$ | $t0$ | $v$ | $t0$ | $v$ | $t0$ | $v$ | $t0$ |
| **OP** | *F* | 4.02 | 3.06 | .283 | 11.27 | .891 | .019 | .002 | 5.07 |
|  | *df* | 1, 19 | 1, 19 | 1, 19 | 1, 19 | 1, 19 | 1, 19 | 1, 19 | 1, 19 |
|  | *p* | .236 | .128 | .801 | **.012** | .714 | .893 | .968 | **.052** |
| | $\eta_p^2$ | .175 | .139 | .015 | .372 | .045 | .001 | .000 | .211 |
| **SOA** | *F* | 2.10 | 1.33 | 28.02 | 43.23 | 7.23 | 9.03 | 3.45 | 5.93 |
|  | *df* | 1.74, 33.18 | 1.44, 27.38 | 3, 57 | 1.40, 26.62 | 3, 57 | 1.73, 32.87 | 3, 57 | 1.93, 36.77 |
|  | *p* | .143 | .319 | **.002** | **.002** | **.002** | **.002** | **.036** | **.008** |
| | $\eta_p^2$ | .100 | .056 | .596 | .695 | .276 | .322 | .157 | .238 |
| **OP x SOA** | *F* | 1.78 | 3.21 | 1.37 | 10.91 | 2.16 | 1.84 | 3.50 | 1.62 |
|  | *df* | 3, 57 | 1.87, 35.55 | 2.09, 39.81 | 3, 57 | 3, 57 | 2.07, 39.43 | 3, 57 | 2.19, 41.60 |
|  | *p* | .188 | .110 | .188 | **.003** | .188 | .208 | .112 | .208 |
| | $\eta_p^2$ | .086 | .145 | .084 | .365 | .100 | .089 | .156 | .079 |

## 4. Discussion

The purpose of this study was to investigate the underlying mechanisms of dual-task interference in a simulated driving environment. We used a systematically controlled dual-task paradigm in which an image task was presented at set times before or after the driving task. We investigated the effect of dual-task, SOA, and unpredictability of task order on subjects’ performance and modeled the results using a DDM. We showed that the RTs of both tasks in the dual-task condition were higher than those in the single task condition. SOA influenced RTs of the two tasks when they were presented second and the RT of the image task when it was presented first. DDM Modeling showed a change in both the drift rate and non-decision times, suggesting that the two tasks were processed in a partial parallel. Unpredictability attenuated the effect of SOA by changing the order of the response to the two tasks. This effect induced a change in the non-decision time of the second task in the DDM.

We observed that in the dual-task conditions, the RTs of both tasks were higher in the dual-compared to the single-task conditions. Similar results have been found in previous studies of dual-task interference in driving (Cooper et al., 2008; Patten, Kircher, Östlund, & Nilsson, 2004; Strayer, Cooper, Turrill, Coleman, & Hopman, 2017; Young, Regan, & Hammer, 2007). We observed this RT difference both for the second and the first task. This observation was inconsistent with Levy et al. (2006) and Hibberd et al. (2013), who showed the interference effect only for the second task. In these two studies, for the driving task, participants performed a car following in which they pressed the brake pedal when the color of the brake light changed. In our study, for performing the lane change, participants had to press a key, hold it, and tune the location of the car to avoid the collision. The more continuous nature of the response in our study might have enhanced the interference effect, causing it to be present even for the first tasks. Another reason for this effect might be that in the dual-task condition, in addition to performing the two tasks, subjects needed to decide about the order of the response to the two tasks. This additional processing might have caused an increase in the RTs (Sigman & Dehaene, 2006). Both Levy et al. (2006) and Hibberd et al. (2013) had instructed the participants to respond according to the presentation order of the stimuli. In our study, however, in order to keep the settings as natural as possible, participants were free to choose the order of their responses. This setting might require additional processing and might contribute to the differences between the dual and single-task conditions.

Similar to many previous studies (Levy et al., 2006; Pashler, 1984; Sigman & Dehaene, 2005), on the second task, we observed a substantial increase in the dual-task effect at short SOAs. This result is predicted by both dual-task models (Pashler, 1994a; Michael Tombu & Jolicœur, 2003). The bottleneck model suggests postponement of the second task is due to the serial processing in the decision stage, and the capacity sharing model suggests that this effect results from resource limitations in the decision process. On the other hand, another finding of our study was that when the image task was presented first, the image RT decreased at shorter SOAs. As mentioned before, the bottleneck model predicts no change in the RT of the first task across SOAs, while the capacity sharing model predicts an increase in the RT of the first task across SOAs. As such, none of these two models are compatible with this part of our results. This effect might be due to the intrinsic time pressure in our driving task that causes participants to try to respond to the image task faster at short SOAs compared to the long SOAs. Overall, based on average RTs alone, it is not possible to choose which one of the two dual-task models best predict the results.

In order to better understand the underlying mechanism of the dual-task effect in driving, we next performed a drift-diffusion analysis. The DDM uncovered that the drift rate of the second task is not constant across SOAs and decreases as the two tasks get closer together in time. This finding suggests that the evidence accumulation for the second task does not stop but continues during the processing of the first task. This result suggests some degree of capacity sharing for the processing of the two tasks. In addition, the t0 of the second task increased with the decrease of the SOA, suggesting some delay in the processing of the second task due to a potential bottleneck. In other words, our results suggest that the best model to account for dual-task interference in driving is a partial parallel model combining the two extremes suggested by capacity sharing and bottleneck models. These results are consistent with the study of Zylberberg et al. (2012), who showed that the decision processes of the two tasks are performed in parallel, however, there is a bottleneck in the mapping of the responses of the two tasks at short SOAs. This bottleneck could also be related to the extra processing required for choosing which task to respond to (Sigman & Dehaene, 2006). It could also be caused by the additional delays imposed by disengagement from the first task (Sigman & Dehaene, 2006). Our current model cannot distinguish the source of this bottleneck. Future studies with more complex models could further illuminate the underlying source of the bottleneck.

The predictability of the order of the tasks had a large effect on the RT of both the image and the driving tasks. This effect was higher in trials in which the driving task was presented first. OP increased the effect of SOA on RT for the first task and attenuated it for the second task. Further analysis showed that the effect of OP was largely caused by the change in the response order of the two tasks. Participants did not always respond to the two tasks according to the presentation order. In general, they had a bias for responding to the driving task first. OP did not change this average bias, but in the unpredictable condition, participants tended to reverse the order of the response in short SOAs, causing a change in the measured dual-task effect. Further, DDM analysis showed that the unpredictability of task order only decreases the non-decision time of the second task, especially at shorter SOAs for the driving task. This effect could have resulted from the change in the response order of the two tasks. The results of this order predictability manipulation suggest that the task presentation order is not the only determinant of the response order, and a top-down task control system might be involved that determines the priority of the two tasks. Our results show that subjects had a bias to respond to the driving task first. This bias might be due to the context of the driving task and the intrinsic time pressure for responding to the driving task in order to avoid collision with the cone obstacles.

Previous studies have also provided evidence in support of a higher-order mechanism to control the response order (De Jong, 1995; Fernández et al., 2011; Leonhard, 2011; Leonhard, Fernández, Ulrich, & Miller, 2011; Luria & Meiran, 2003; Ruiz Fernández et al., 2013; Sigman & Dehaene, 2006; Szameitat et al., 2006). Based on the optimization model of Miller et al. (2009), the participants’ aim in a dual-task paradigm is to decrease the total RTs (RT of the first task + RT of the second task). According to this model, the participants respond to the easy task sooner than the difficult one. In other words, the duration of the components of the two tasks determines which task is responded to first. A few studies have confirmed this prediction. For example, Sigman and Deheane (2006) have shown that when task order is unpredictable, the task with shorter perceptual duration is responded to first. In another study, Ruiz Fernández et al. (2011, 2012) showed that the duration of the decision and the motor stages could influence the order of the response to the two tasks. The result of our study is not fully compatible with the predictions of the optimization model. Although the motor stage of the driving task was difficult and longer than the image task, participants prioritized the driving over the image task. In contrast to our study, all previous studies that provided evidence for the optimization model have used artificial dual-task paradigms. Thus, it appears that in the real world dual-task conditions, additional factors also influence the priority to respond to the tasks. For example, in our study, there is an intrinsic time pressure for responding to the driving task. To avoid a collision, participants should change the driving lane as soon as possible. This time pressure might be the underlying reason for the increased priority of the driving task.

## 5. Conclusions

To sum up, we observed a strong effect of dual-task interference in driving. This effect is modulated by SOA and is attenuated by the predictability of the task order. Furthermore, DDM showed that the two tasks are processed in a partially parallel manner. Our results could be applicable for optimizing the timing of driving assistance systems such as the road signs, alarm systems, and other driver interfaces in order to reduce accidents.

## Acknawledgements

We thank Mahdi Shafiei for his assistance in developing the simulated driving environment and Sajjad Zabbah for his assistance in drift diffusion modeling. This research was supported in part by the Intramural Research Program of the NIH, National Institute of Mental Health (ZIA MH002035-39). The collection and analysis of the data was performed entirely at the Institute for Research in Fundamental Sciences (IPM) supported by a research grant from IPM.

